# Phylogeny of the Damselfishes (Pomacentridae) and Patterns of Asymmetrical Diversification in Body Size and Feeding Ecology

**DOI:** 10.1101/2021.02.07.430149

**Authors:** Charlene L. McCord, Chloe M. Nash, W. James Cooper, Mark W. Westneat

**Affiliations:** California State University Dominguez Hills, College of Natural and Behavioral Sciences, 1000 E. Victoria Street, Carson, CA 90747; Western Washington University, Department of Biology and Program in Marine and Coastal Science, 516 High Street, Bellingham, WA 98225; University of Chicago, Department of Organismal Biology and Anatomy, and Committee on Evolutionary Biology, 1027 E. 57^th^ St, Chicago IL, 60637, USA; Field Museum of Natural History, Division of Fishes, 1400 S. Lake Shore Dr., Chicago, IL 60605

**Keywords:** Pomacentridae, phylogenetics, body size, diversification, evolution, ecotype

## Abstract

The damselfishes (family Pomacentridae) inhabit near-shore communities in tropical and temperature oceans as one of the major lineages in coral reef fish assemblages. Our understanding of their evolutionary ecology, morphology and function has often been advanced by increasingly detailed and accurate molecular phylogenies. Here we present the next stage of multi-locus, molecular phylogenetics for the group based on analysis of 12 nuclear and mitochondrial gene sequences from 345 of the 422 damselfishes. The resulting well-resolved phylogeny helps to address several important questions about higher-level damselfish relationships, their evolutionary history and patterns of divergence. A time-calibrated phylogenetic tree yields a root age for the family of 55.5 mya, refines the age of origin for a number of diverse genera, and shows that ecological changes during the Eocene-Oligocene transition provided opportunities for damselfish diversification. We explored the idea that body size extremes have evolved repeatedly among the Pomacentridae, and demonstrate that large and small body sizes have evolved independently at least 40 times and with asymmetric rates of transition among size classes. We tested the hypothesis that transitions among dietary ecotypes (benthic herbivory, pelagic planktivory and intermediate omnivory) are asymmetric, with higher transition rates from intermediate omnivory to either planktivory or herbivory. Using multistate hidden-state speciation and extinction models, we found that both body size and dietary ecotype are significantly associated with patterns of diversification across the damselfishes, and that the highest rates of net diversification are associated with medium body size and pelagic planktivory. We also conclude that the pattern of evolutionary diversification in feeding ecology, with frequent and asymmetrical transitions between feeding ecotypes, is largely restricted to the subfamily Pomacentrinae in the Indo-West Pacific. Trait diversification patterns for damselfishes across a fully resolved phylogeny challenge many recent general conclusions about the evolution of reef fishes.

## Introduction

Advances in the richness and accuracy of the Tree of Life are critically important for understanding broad evolutionary patterns and processes, as well as exploring the unique features of specific groups of organisms that capture our attention. It is a central challenge of systematics to resolve phylogenies that include large numbers of species and include previously unexamined taxa in well-supported trees, thus improving our ability to test evolutionary hypotheses and examine patterns of diversification. In recent years, we have gained an increasingly clear picture of the phylogenetic relationships among coral reef fishes through the pursuit of densely sampled species phylogenies for diverse reef fish families (1–13). By combining species-rich phylogenies with rich data layers of functional and ecological traits, significant advances have been made in our understanding of the diversification of fish lineages in coral reef ecosystems (14–22). The project presented here aims to enhance the species richness present in the phylogeny of the reef fish family Pomacentridae and examine the evolutionary history of key structural and ecological traits of the damselfishes, with a focus on the transitions in body size and dietary ecotype throughout pomacentrid diversification.

The damselfishes (Pomacentridae) are a major component of the global coral reef and temperate rocky reef ichthyofauna (23,24) and their systematic study has an unusually deep and rich history. Descriptions of damselfishes are found in publications that laid the foundations of taxonomy such as Linnæus’s Systema Naturae (25), the beginnings of ichthyology with Artedi’s *Ichthyologia* (26), and even the entirety of western biology with Artistotle’s *History of Animals*, (27). Damselfishes were studied by several of the most prominent naturalists of the 19^th^ century including Lacépède (28) and Cuvier (29), and the type specimen of one damselfish genus (*Stegastes*) was collected by Charles Darwin early in the voyage of *H*.*M*.*S. Beagle* (30). The past 60 years have produced a massive surge in marine research, including the publication of several hundred studies of pomacentrid fishes. Advances in molecular biology and computational ability have facilitated increasingly comprehensive molecular phylogenetic studies of damselfish relationships (1,15,31–37). These phylogenies have been used in multiple phylogenetic comparative studies that have examined the evolution of their functional morphology and ecological diversity (14,15,38–41).

The Pomacentridae range in size over an order of magnitude from the small *Chrysiptera giti* (4.5 cm) up to the giant *Microspathodon dorsalis* (45 cm), and occupy marine habitats from shallow coastal waters down to 200m depths. The circumtropical distribution of damselfishes is centered on marine coral and rocky reef habitats, with a variety of temperate species occurring at up to 50° North and South latitude. Members of the family are known for intriguing behaviors such as strong territorial aggression, and complex farming or gardening behaviors in which dense stands of filamentous algae are tended for food (42). Damselfishes have been placed in broad ecological-functional categories (“ecotypes”) using combinations of traits such as diet (planktivory, herbivory, and omnivory) and primary feeding location in the water column (benthic, pelagic, intermediate) that are useful for exploring the evolution of damselfish ecomorphology (14,15). Detailed analyses of functional morphology have shown that mandible morphology and jaw protrusion ability are strong predictors of damselfish ecotype (14,15,39) and that trophic specialization is tightly tied to habitat and social behavior (38). Evolutionary-developmental studies of damselfishes have also provided insight into the ontogenetic changes associated with shifts in adult feeding ecology (43–46).

A central goal of this study is to explore traits associated with diversification across the pomacentrid phylogeny. A noteworthy evolutionary pattern among damselfishes is that dietary ecotype has been shown to be unevenly distributed across the Pomacentridae, with transition rates from the intermediate to specialized dietary ecotypes shown to be higher than in the opposite direction (38). Similarly, body size has been shown to be highly variable across damselfishes, although evolutionary patterns of size have received less attention (14, 44). There are frequent switches back and forth between ecotype states and size categories within some damselfish groups, creating a strong pattern of convergence (14), although there is a trend in some groups to become fixed on one of the specialist ecotypes, creating asymmetrical patterns of diversification among clades. However, previous work has not found strong evidence that differences in diversification rates among feeding strategies support an evolutionary “dead-end” hypothesis, in which specialization results in reduced speciation and elevated extinction rates (47,48). Given that the asymmetry in state distribution could be the result of variation in transition rates or differences in diversification rates associated with each state, it is important to jointly estimate these two processes (49). Recent advances in the hidden-state speciation and extinction modeling framework allow us to examine this variation in diversification rates and its potential association with body size and dietary ecotype using multistate characters (50). Therefore, to build on recent work in this area (38) and explore the role of trait transition asymmetry in damselfish diversification, we tested these hypotheses using multistate hidden state modeling, using a robust phylogeny and trait database composed of 345 species.

In this study we present advances in our understanding of damselfish phylogenetic relationships based on molecular sequence data for 12 loci from 345 of the 422 extant pomacentrid species (51). This analysis is compared to prior work on about 210 species and the contemporaneous work by Tang et al. (37) using 8 loci from 322 damselfish species. We use the time-calibrated phylogeny to examine evolutionary rates of transition across several important ecological traits to test hypotheses about the complex relationship between trait evolution and the tempo of damselfish diversification, and examine the origins of major clades across space and time. Our central aims were to generate a new species-rich phylogenetic framework, use that framework to provide new estimates for the divergence times of multiple damselfish clades, reconstruct the evolution of pomacentrid body size and feeding ecology, and explore the evolutionary patterns associated with the strikingly asymmetrical distributions of body size among clades and of benthic, pelagic and intermediate ecotypes across the damselfish phylogeny.

## Methods

### Species sampling and genes sequenced

DNA sequence data for 350 fish species were analyzed in this study, including 345 species of damselfishes and 5 outgroup taxa from two additional families in the Ovalentaria: Embiotocidae and Cichlidae (See Table S1 for a list of all taxa and genes examined, with large data sets contributed by past authors color coded to provide clear attribution). At least one member of each damselfish genus was included in our phylogenetic analyses. Specimens sequenced for the present study (180 sequences for 80 species, including 10 not previously sequenced) were collected by the authors, purchased at local fish markets, or borrowed from outside institutions and most are associated with a voucher specimen (Table S1). We analyzed portions of twelve genes totaling 8238 bases from seven mitochondrial regions: 12s (901 bp), 16s (559 bp), ND3 (431 bp), ATP (842 bp), COI (651 bp), Cytb (1140 bp), and D-loop (439); and five nuclear loci: RAG1 (900 bp), RAG2 (800 bp), DLX2 (482 bp), Tmo4C4 (417 bp), and BMP4 (548 bp). Most nuclear sequences and three mitochondrial regions (12s, 16s and ND3) were sequenced at the Pritzker Laboratory of the Field Museum of Natural History. For new gene sequences, DNA sequencing protocols were similar to those described in previous phylogenetics studies from our laboratory (1,12,13), see S1Text, supplemental methods. Sequence data for four additional mitochondrial loci (ATP, COI, CytB and D-loop) were downloaded from GenBank and included in the concatenated supermatrix (Table S1). Taxonomic names in the present study follow those proposed by Tang et al. (37) after we released our work on BioRxiv (52), and we note that our project includes the clarified taxonomic issues and resolved sequence identifications from this work. Important differences between the two studies in data set construction and sequence alignment are included in the S1Text supplemental methods.

### Phylogenetic analysis

The best fit models of nucleotide evolution for damselfish gene sequences were selected using PartitionFinder2 (53) and maximum likelihood analyses were performed using Garli2 (54). An initial set of eight parallel (mpi) analyses (three replicates each; 24 runs) was performed using a random starting tree. The topology with the highest likelihood from this first run was used as the starting tree for a second set of 24 simultaneous maximum likelihood analyses. Termination condition settings for all analyses were (1) number of generations without topology improvement (genthreshfortopoterm) = 50000, (2) max score improvement over recent generations required for termination (scorethreshforterm) = 0.05 and required score improvement for topology to be considered better (significanttopochange) = 0.01.

Bayesian phylogenetic analyses of the concatenated DNA supermatrix were conducted using MrBayes (55) on an Asus GL12CM gaming PC (Intel i7-8700K, NVIDIA GTX 1080 GPU). We performed two independent runs of a Markov Chain Monte Carlo analysis with four chains (eight chains total) running in parallel for 50 million generations. Trees were saved every 1000 generations for each run and used default priors for the transition/transversion rate ratio, branch length, alpha parameter of the gamma distribution for rate heterogeneity, proportion of invariant sites, base frequencies and tree topology parameters. To help ensure that stationarity was reached, the first 10 million generations (10000 sampled trees) were discarded from each run as burn-in and the remaining 40 million generations (45000 sampled trees for each of the four runs) were used in subsequent analysis. To determine the posterior probability of clades, a majority-rule consensus tree calculated from the post burn-in trees was constructed. The majority-rule consensus tree and maximum posterior probability phylogenetic trees were then exported and viewed using FigTree version 1.3.1 (56) and the ggtree package (57) in R version 3.3.1 (R Core Team).

### Divergence time estimation

To explore the timing of damselfish divergence events we calibrated two nodes in the pomacentrid tree using fossil evidence. *Palaeopomacentrus orphae* and *Lorenzichthys olihan* from the Ypresian were used to determine the minimum age of the crown pomacentrids (58). *Chromis savornini* from the late Miocene 6.5 Ma (59) was used to date the origin of the genus *Chromis*. Due to the likely young age of *Chromis savornini*, we assigned a soft upper bound to this calibration point (48.5 Ma, age of the oldest crown pomacentrid). Outside the family Pomacentridae, we also set a prior age estimate of 96 mya for the most recent common ancestor of cichlids, embiotocids, and pomacentrids within the Ovalenteria (60). To estimate divergence times, we used a relaxed clock of log normal distributed rates in BEAST 2.6.3 (61). A standard birth-death rate model was used to estimate rates of cladogenesis. We partitioned the twelve genes used in these analyses such that each region was modeled separately under its best-fit model of substitution. Tracer v1.6 was used to assess convergence following two independent analyses of 100 million generations each. The first 25% of generations were used as burn-in and the effective sample size (ESS) for all model parameters were checked for good mixing of the final Markov Chain Monte Carlo (all ESS exceeded 200 in all analyses).

### Damselfish trait matrix assembly and analysis

Data for body size (body length), body depth (as % body length), dietary ecotype (benthic, pelagic, or intermediate), and presence of algal farming behavior were gathered from prior work (14,15), original species descriptions, or mined from FishBase using the rFishBase package in R (62). Data gathered from repositories were checked against original literature whenever possible (Table S3, Table S4). Continuous data for total body length were binned into three discrete states using gap coding (63). Size classes in the 3-state character were defined as small (S = 4.5-10.2 cm TL), medium (M = 10.5-14.1 cm TL) and large (L = 14.4-45cm TL). Using the ‘ace’ function in the R package ape (64), ancestral character states were estimated across the phylogeny using the transition rate models all rates different (ARD), symmetrical transitions (SYM), and equal rates (ER). The ARD model was selected for all traits in order to plot character trait maps. The number of transitions between states for body size and ecotype were computed using the countsimmap function in the R package Phytools (64). Lineage through time plots were performed using the ltt.plot function in ape.

We also tested the hypothesis that simultaneous loss and gain of a complex trait, such as body size or dietary ecotype, without passing through an intermediate state, may be rare or impossible by constructing a set of “constrained” ace transition models. This idea is supported by previous work which found no direct transitions between the benthic and pelagic states (38). These “constrained” state transition models require transitions through the intermediate state.

Tip states were mapped onto the time-calibrated phylogeny, and the estimated likelihoods of each ancestral state were mapped onto internal nodes within the phylogeny, allowing visualization of independent trait origins and convergence across clades. We tested for strength of phylogenetic signal for continuous traits (body size, body depth) by computing Blomberg’s K and Pagel’s lambda using the ‘phylosig’ function in the R package Phytools (64). Statistical support was computed using 1000 randomizations. For the discrete traits of body size and ecotype, we conducted discrete lambda tests using likelihood (lambda = 0 vs estimated lambda) using the ‘fitDiscrete’ function in the R package Geiger (65). To test for the association of dietary ecotype and farming behavior with size, we also examined trait associations across the phylogeny using a phylogenetic ANOVA with 10,000 simulations, computed using the ‘phylANOVA’ function in Phytools.

### Analysis of Ecotype Dependent Diversification using MuHiSSE

We used a multistate, hidden-state speciation and extinction (MuHiSSE) model framework (66) to evaluate the association of transitions among body size categories and dietary ecotypes with rates of diversification across the damselfish phylogeny. For this analysis, each discrete ecotype was coded as the presence or absence of two binary states. For size the trait was coded as small (01), large (10) or medium (11), and ecotype was coded as benthic (01), pelagic (10) or intermediate (11). Because no direct transitions between small and large or between pelagic and benthic are found, and the constrained transition models have the highest likelihood, we used the “constrained” transition model for all SSE analyses. We designed 8 hidden state models (see TextS1 supplemental methods for details) to test specific hypotheses about the influence of size and ecotype on rates of diversification and explore the level of asymmetry in transitions associated with size and ecotype. These models fall into five distinct categories: 1) “trivial null” model, which assumes a single rate of diversification across phylogeny independent of character state, 2) state dependent diversification, which assumes diversification parameters are shared among lineages with the same ecotype (MuSSE), 3) state dependent diversification with hidden state(s), in which the diversification rates are shared among ecotype in combination with a hidden (non-focal) state (MuHiSSE), 4) state absorbing, in which transitions out of a particular state were constrained to zero (i.e. Benthic → Intermediate = 0) and 5) character independent diversification, which assumes diversification parameters are independent of ecotype but allowed to vary over increasing numbers of hidden states (MuCID models; Table 1). Given the prevalence of recovering a “false positive” when using SSE models, MuCID models can act as the proper null model due to the partitioning of rate variation across hidden states that are not attributable to the focal state (66,67).

**Table 1.** Ranked MuHiSSE results for body size and ecotype, in descending order of support for tested models.

| Body Size Model | Brief Description | Free Parameters | Likelihood | AIC | ΔAIC | ωAIC |
| --- | --- | --- | --- | --- | --- | --- |
| MuHiSSE Relaxed | Relaxed state dependent diversification with a hidden state | 26 | -1414.42 | 2875.98 | 0.00 | 0.57 |
| Large Absorbing | No transitions out of large state | 21 | -1417.00 | 2876.59 | 0.60 | 0.42 |
| MuHiSSE | State dependent diversification with a hidden state | 25 | -1419.30 | 2883.46 | 7.48 | 0.01 |
| Small Absorbing | No transitions out of small state | 21 | -1428.61 | 2899.82 | 23.84 | 0.00 |
| CID2 | Character-independent with 2 hidden states | 17 | -1441.98 | 2904.62 | 28.63 | 0.00 |
| MuSSE | State dependent diversification | 12 | -1442.17 | 2909.28 | 33.30 | 0.00 |
| "Dull" | Equal rates | 7 | -1447.66 | 2911.75 | 35.77 | 0.00 |
| CID3 | Character-independent with 3 hidden states | 19 | -1440.25 | 2914.16 | 38.17 | 0.00 |

| Ecotype Model | Brief Description | Free Parameters | Likelihood | AIC | ΔAIC | ωAIC |
| --- | --- | --- | --- | --- | --- | --- |
| MuHiSSE | State dependent diversification with a hidden state | 25 | -1354.08 | 2753.01 | 0.00 | 0.46 |
| MuHiSSE Relaxed | Relaxed state dependent diversification with a hidden state | 26 | -1353.20 | 2753.55 | 0.53 | 0.35 |
| Benthic Absorbing | No transitions out of benthic state | 21 | -1356.15 | 2754.88 | 1.87 | 0.18 |
| CID2 | Character-independent with 2 hidden states | 17 | -1377.39 | 2775.44 | 22.43 | 0.00 |
| CID3 | Character-independent with 3 hidden states | 19 | -1375.13 | 2783.91 | 30.90 | 0.00 |
| Pelagic Absorbing | No transitions out of pelagic state | 21 | -1372.04 | 2786.66 | 33.65 | 0.00 |
| MuSSE | State dependent diversification | 12 | -1498.18 | 3017.02 | 264.00 | 0.00 |
| "Dull" | Equal rates | 7 | -1504.26 | 3020.77 | 267.76 | 0.00 |

Net diversification (defined as speciation divided by extinction), extinction fraction (extinction/speciation) and transitions among states were either constrained or allowed to vary depending on the model (see S1Text). Although there are known issues in estimating extinction in extant only phylogenies, our models allowed it to vary in the same way as speciation. The reparameterization of net turnover and extinction fraction alleviates problems associated with over-fitting when speciation and extinction are highly correlated, but both matter in explaining the diversity pattern (67). To account for sampling bias among character states, we used the following sampling fraction, which corresponds to the percentage of taxa in our tree with a particular state over the total number of taxa with that trait: Pelagic = 0.68, Benthic = 0.72, Intermediate = 0.76.

Following Nakov et al. (50) we used 100 random starting points to account for the initial run not recovering the MLE for the optimization of each model. Models were optimized using the MuHiSSE function in the hisse package in R (66,68) on the midway2 computing cluster through the Research Computing Center at the University of Chicago. The parameters from the optimization iteration with the highest likelihood for each model were ranked according to AICc. Diversification and transition parameters were estimated using a model averaging approach through weighted AIC (ωAIC), which reduces the subjectivity of choosing the thresholds for model choice and accounts for the uncertainty around parameter estimates across hidden states (69). This was done by averaging each estimated rate of diversification per-lineage using the weighted probability of occupying an alternative hidden state (50). We reconstructed the marginal ancestral states according to ωAIC using the MarginReconMuHiSSE function and approximated the likelihood surface around the maximum likelihood estimates with an adaptive sampling method using the SupportRegionMuHiSSE function, in the hisse package (66) and figures were made using the utilhisse package in R (50). All R scripts are available at https://github.com/cmnash/pomacentridae_supp_code.

## Results

Phylogenetic analyses revealed a strongly supported monophyletic Pomacentridae that originated in the Lower Eocene (55.5 mya) with the five modern subfamilies (70) arising during the mid-Eocene to upper Oligocene, from about 50-30 mya (Fig. 1 and 2). Major clades of damselfishes were recovered with high support and topologies similar to those reported in prior studies, yet with some significant revisions of relationships within several clades. A central result of this study is that deep branching events within Pomacentridae were followed by a remarkably steady rate of clade diversification over the last 50 million years. Body size shows weak phylogenetic signal, with multiple instances of convergence on both large and small body size across the phylogeny. Repeated patterns of evolutionary change in both body size and feeding ecology across the phylogeny show that patterns of species diversification are associated with an imbalanced or asymmetric frequency of transition between states. A frequent pattern of back-and-forth transitions among ecotypes is primarily restricted to the crown group Pomacentrinae.

**Figure 1.**
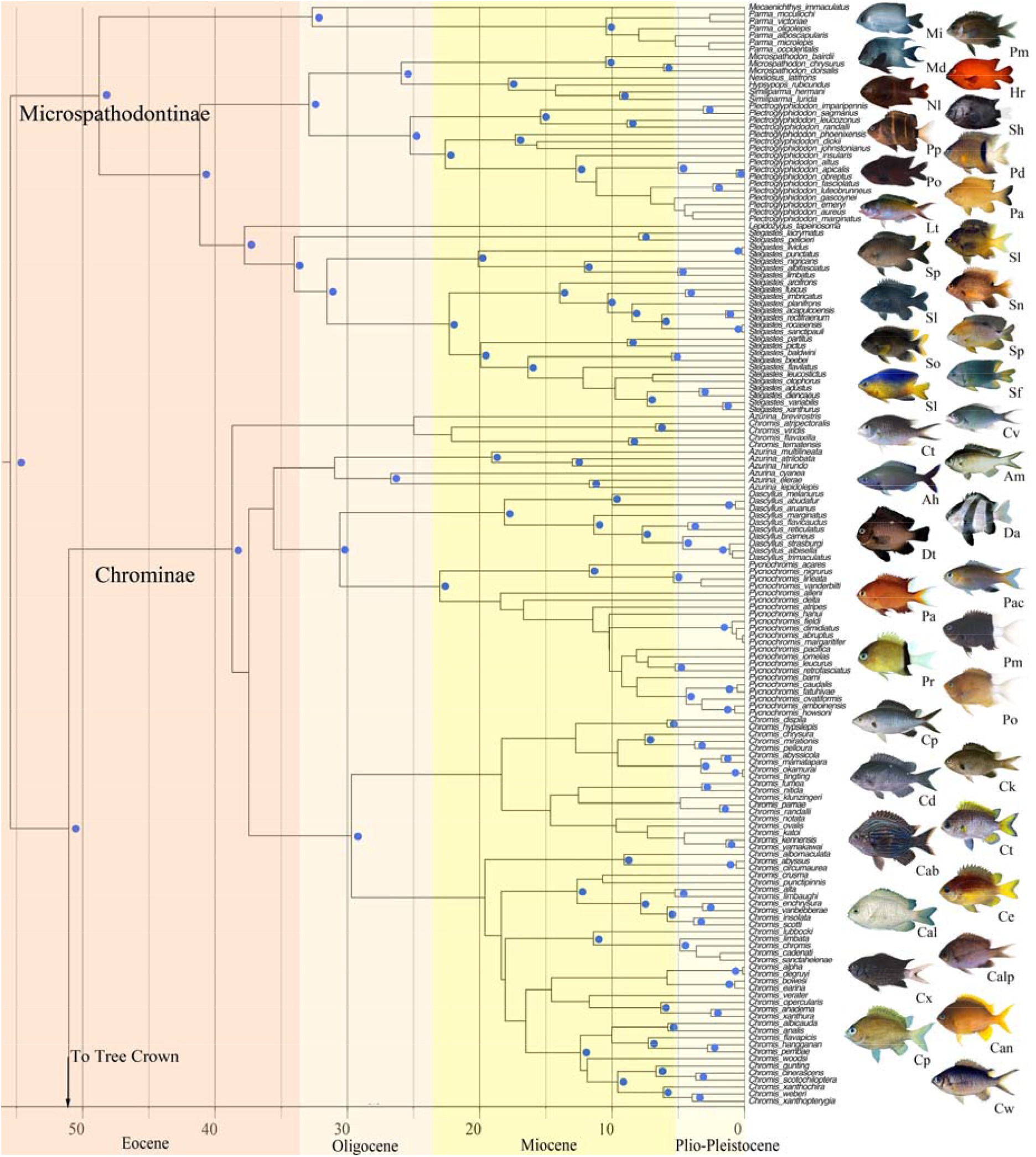
Time-calibrated phylogenetic tree of the Pomacentridae. Lower half of the tree including the subfamilies Microspathodontinae and Chrominae. Time axis in millions of years before present. Nodal values with Bayesian posterior support levels above 0.9 are indicated with blue dots (see Fig. S2 for all support values and Fig. S3 for all ages). Representative photos are labeled with the first letter of genus and species, matching species names in the tree closest to the photo.

**Figure 2.**
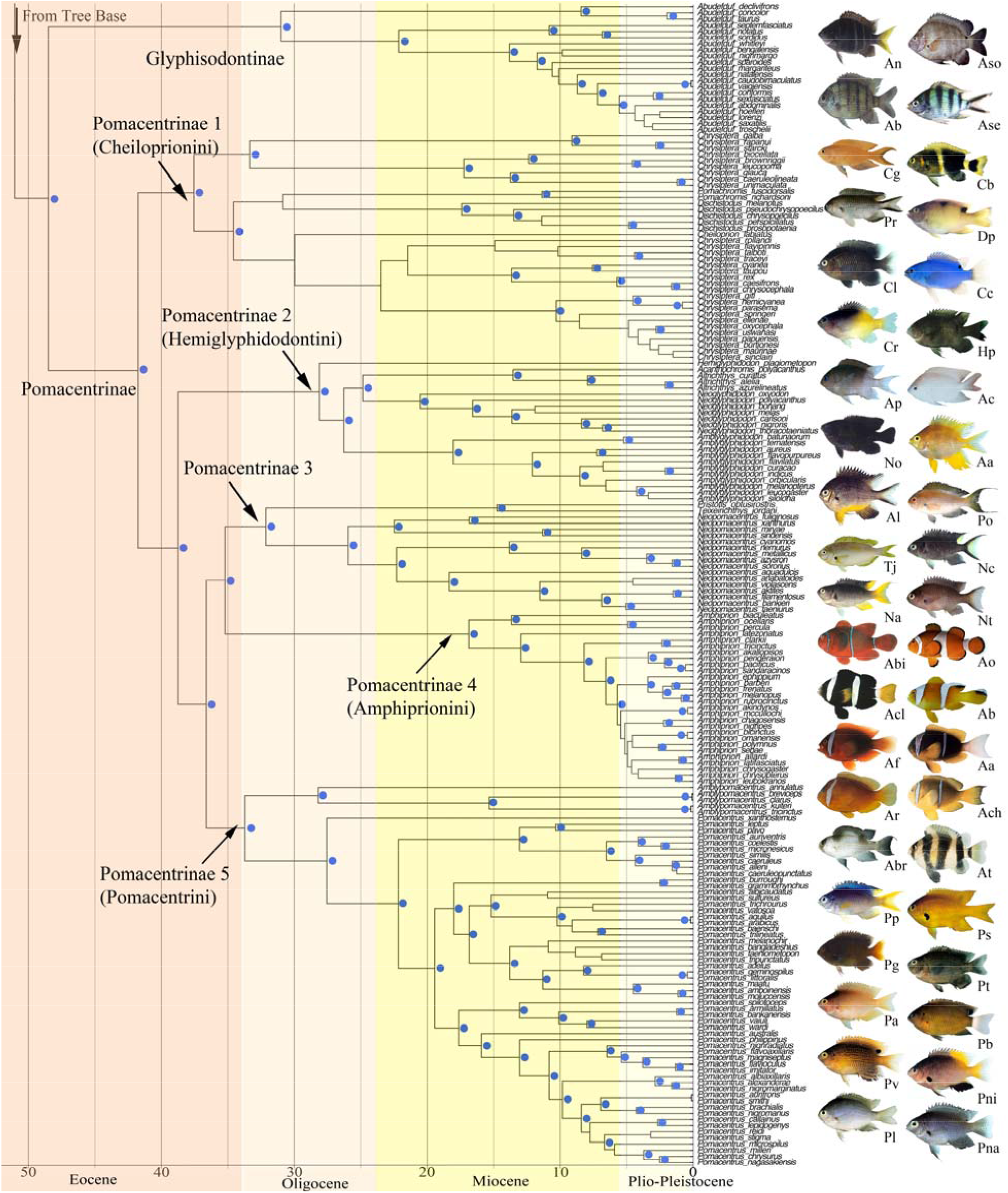
Time-calibrated phylogenetic tree of the Pomacentridae. Upper (crown) half of the tree including the subfamilies Glyphysodontinae and Pomacentrinae. Time axis in millions of years before present. Nodal values with Bayesian posterior support levels above 0.9 are indicated with blue dots (see Fig. S2 for all support values and Fig. S3 for all ages). Representative photos are labeled with the first letter of genus and species, matching species names in the tree closest to the photo.

### Pomacentrid phylogenetics

Phylogenetic analyses resolved a monophyletic Pomacentridae with four well-resolved and strongly supported damselfish subfamilies (Fig. 1, 2), supporting recent subfamily taxonomy of the family (37,70). The same overall damselfish topology and subfamily structure are well-supported by both ML and Bayesian analyses (S1Fig, S2Fig). The sister group to the rest of the damselfishes is the subfamily Microspathodontinae, which comprises three well-supported (*pp =* 1.0) clades. The sister-group to the rest of the Microspathodontinae contains the monotypic genus *Mecaenichthys* and a monophyletic group of four *Parma* species present in the tree (Fig. 1). Moving upward from the base, the second major clade of the Microspathodontinae contains the giant damselfishes, a clade composed of the genera *Microspathadon, Nexilosus, Hypsypops* and *Similiparma* (70). *Nexilosus* is represented by COI barcode sequence only, and its position is often unstable within this clade. The giant damsels are sister to the genus *Plectroglyphidodon*, which is now monophyletic (Fig. 1) after several *Stegastes* were recently renamed (37). *Lepidozygus tapeinosoma* is sister to the third major branch of the subfamily, *Stegastes* (Fig. 1), with the longest monotypic branch in our trees (Fig. 1). The genus *Stegastes* is well supported as a monophyletic group with new taxonomic names (37) and is an area of high topological support among species (Fig. 1).

The damselfish subfamily Chrominae contains four genera (*Azurina, <u>Chromis</u>, Dascyllus* and *Pycnochromis* (37)) forming several well-resolved major clades, and others with low support (Fig. 1). The sister group to the rest of Chrominae, (Fig. 1: Chrominae1) is a small *Chromis* clade composed of *Chromis flavaxilla, C. ternatensis, C. atripectoralis* and *C. viridis*. Moving up the chromine backbone is the weakly supported *Azurina* clade (Fig. 1), and the strongly supported monophyletic genus *Dascyllus* (Fig. 1). *Dascyllus* forms the sister group to the chromine clade of 22 species that have been moved from *Chromis* to *Pycnochromis* (37). The largest clade of Chrominae is strongly supported as monophyletic and contains 55 of the *Chromis* species we examined (Fig. 1). The phylogeny is split into two main groups, neither of which has strong support nor particularly well-resolved internal relationships, although several tip clades are well supported. Many recently described species of *Chromis*, often with COI barcodes as the only source of data currently available, are in this group.

The monophyletic *Abudefduf* is the next major clade up the damselfish backbone, with high posterior probability (1.0). This is the subfamily Glyphisodontinae, with *Abudefduf declivifrons, A. concolor* and *A. taurus* sister to the rest of the clade (Fig. 2). Moving distally from the base of *Abudefduf*, the next clade is composed of *A. conformis, A. septemfasciatus, A. notatus* and *A. sordidus*. A third clade forms a nearly ladder-like topology of 15 species from the basal node of *A. whitleyi* to the crown sister-pair of *A. hoefleri* and *A. lorenzi* (Fig. 2).

The Pomacentrinae form the largest damselfish subfamily (Fig. 2). This strongly supported clade of 16 genera contains 218 species, more than half of the known damselfishes (51) representing the most diverse lineage in the family. The Pomacentrinae contains the genera *Acanthochromis, Altrichthys, Amblygliphidodon, Amblypomacentrus, Amphiprion, Cheiloprion, Chrysiptera, Dischistodus, Hemiglyphidodon, Neoglyphidodon, Neopomacentrus, Pomacentrus, Pomachromis, Premnas, Pristotis* and *Teixeirichthy*. These taxa are resolved into five major clades. With the recent phylogenetically informed taxonomic changes (37), all genera are either monotypic or monophyletic, with the exception of *Chrysiptera* (Fig. 2).

The sister-clade to the rest of the subfamily (Pomacentrinae 1: tribe Cheiloprionini) is the assemblage of *Chrysiptera* species with two lineages of species from other genera nested among them. One of these contains the genera *Pomachromis* and *Dischistodus* as two monophyletic sister groups. The other is composed of *Cheiloprion labiatus*, an unusual coral polyp specialist that is the only member of its genus. The second pomacentrine lineage (Pomacentrinae 2: tribe Hemiglyphidontini) is composed of five monophyletic genera. The monotypic *Hemiglyphidodon* (*H. plagiometopon*) branches from the basal node, and the next lineage is the monophyletic genus *Amblyglyphidon*, followed by *Acanthochromis* (*A. polyacanthus*) + *Altrichthys* (monophyletic), and the monophyletic *Neoglyphidodon*. These relationships have strong support with just a few nodes of low support within genera.

Pomacentrinae 3 and 4 (Fig. 2) are sister clades, with low support, with group 3 (not presently given tribal status) containing *Pristotis, Teixeirichthys and Neopomacentrus*, and group 4 the monophyletic tribe Amphiprionini. Relationships within Pomacentrinae 3 are strongly supported, with the sister-pair *Pristotis obtusirostris* + *Teixeirichthys jordani* forming the sister lineage to *Neopomacentrus*. Pomacentrinae 4 (Fig. 2) is the anemonefish tribe Amphiprionini.

Pomacentrinae 5 (Fig. 2) contains the monophyletic genus *Pomacentrus* plus *Amblypomacentrus*, which now includes three problematic *Chrysiptera* species that have been reassigned to *Amblypomacentrus* (37). These relationships are all well-supported. The *Amblypomacentrus* lineage is sister to a monophyletic *Pomacentrus* clade that contains 57 species of the 77 presently known (51). Except for several weak nodes out toward the tips, the topology of the *Pomacentrus* phylogeny inferred here is well-resolved. *Pomacentrus xanthosternus* is resolved as sister to the rest of the genus, with successive clades suggesting likely group memberships for many recently described *Pomacentrus* represented by small amounts of sequence data.

### Dates of origin and divergence times for damselfishes

Time-calibrated phylogenetic analysis of the present data set (see Fig. S3 for all node ages) suggests that the damselfishes split from embiotocids approximately 80 mya, with the root node of the first family divergence for the Pomacentridae at 55.5 mya. The Microspathodontinae diverged from other pomacentrids at this time and underwent important vicariance events at 49, 41, 38, and 32 mya (Fig. 1). The tribe Microspathodontini (the giant damselfishes) began to diversify ∼26 mya (Fig. 1). The Chrominae originated 51 mya and extant lineages began to diverge ∼38 mya (Fig. 1). Divergence dates for major Chrominae clades range from 39 to about 30 mya. *Dascyllus* arose ∼31 mya and existing clades began to diversify ∼18.1 mya (Fig. 1). *Pycnochromis* began to radiate earlier than *Dascyllus* (about 23 mya). *Chromis* bifurcated ∼30 mya and continued to diversify at a relatively constant rate throughout the Miocene. Most of the *Chromis* species we examined arose during the past 5-15 million years (Fig. 2). The Glyphisodontinae (*Abudefduf*) split from the Pomacentrinae ∼49 mya and the radiation of living *Abudefduf* began in the late Oligocene (∼31 mya).

The initial divergence of the Pomacentrinae and origin of the Pomacentrinae 1 clade occurred ∼41.7 mya (Fig. 2), with group 1 diversifying at a relatively steady rate since its origin. Pomacentrinae 2 and 5 arose 36-37 mya, and Pomacentrinae 3 and 4 diverged from each other ∼35 mya. The *Pomachromis-Dischistodus* clade arose within Pomacentrinae ∼31 mya, and *Cheiloprion* split within Pomacentrinae 1 ∼29 mya. (Fig. 2). Within Pomacentrinae 2 the major clades diverged at the following estimated times: *Hemiglyphidodon* (28 mya); *Amblyglyphidodon* (26 mya); *Acanthachromis* + *Altrichthys* (arose 24 mya; diverged into monophyletic genera 10 mya); and *Neoglyphidodon* (24 mya).

Pomacentrinae 3 (*Neopomacentrus, Pristotis* and *Teixirichthys*) split from Pomacentrinae 4 (the Amphiprionini) ∼35 mya (Fig. 2). This clade diverged steadily over the past 25 my, with the Amphiprionini being relatively young, having diverged from a common ancestor ∼18 mya and most species arising only 3-5 mya. Pomacentrinae 5 diverged ∼34 mya, with diversification of *Pomacentrus* into 77 known species over the past ∼27 million years accounting for nearly one fifth of the damselfishes and a large portion of the Indo-West Pacific coral-reef fish fauna.

### Trait evolution in damselfishes

Ancestral state estimation of body size (Fig. 3) shows that large body size has evolved repeatedly, up to 44 times, throughout the tree (from medium-sized relatives) and that small body size has undergone a similar number (37) of transitions from medium size across the topology. Patterns of transition between body size states are strongly asymmetrical (Fig. 3), with all transitions going through the intermediate state, and transitions directly between small and large size states absent. The giant damselfishes of the tribe Microspathodontini and the Glyphisodontinae (genus *Abudefduf*) exhibit concentrations of large species, while the various lineages of *Chromis, Chrysiptera* and *Pomacentrus* show a preponderance of small species. The continuous trait for body length, the discretized 3-state character for body length shown in Fig. 3, and body depth (as a proportion of length) all show low phylogenetic signal, significantly different from the expectation of Brownian motion across the damselfish tree. For body length, Blomberg’s K was 0.29 (p = 0.001) and Pagel’s lambda was 0.86 (p < 0.001) reflecting the tendency for some clades to have a particular size trait, but others showing a distribution of small, medium and large taxa. Body depth ranged from about 27% SL in slender species such as *Azurina* to 66% in deeper bodied forms such as *Amblyglyphidodon*, with K= 0.16 (p = 0.001) and λ=0.87 (p < 0.001). This indicates that close relatives were significantly more variable in both body depth and length than would be expected under Brownian motion.

**Figure 3.**
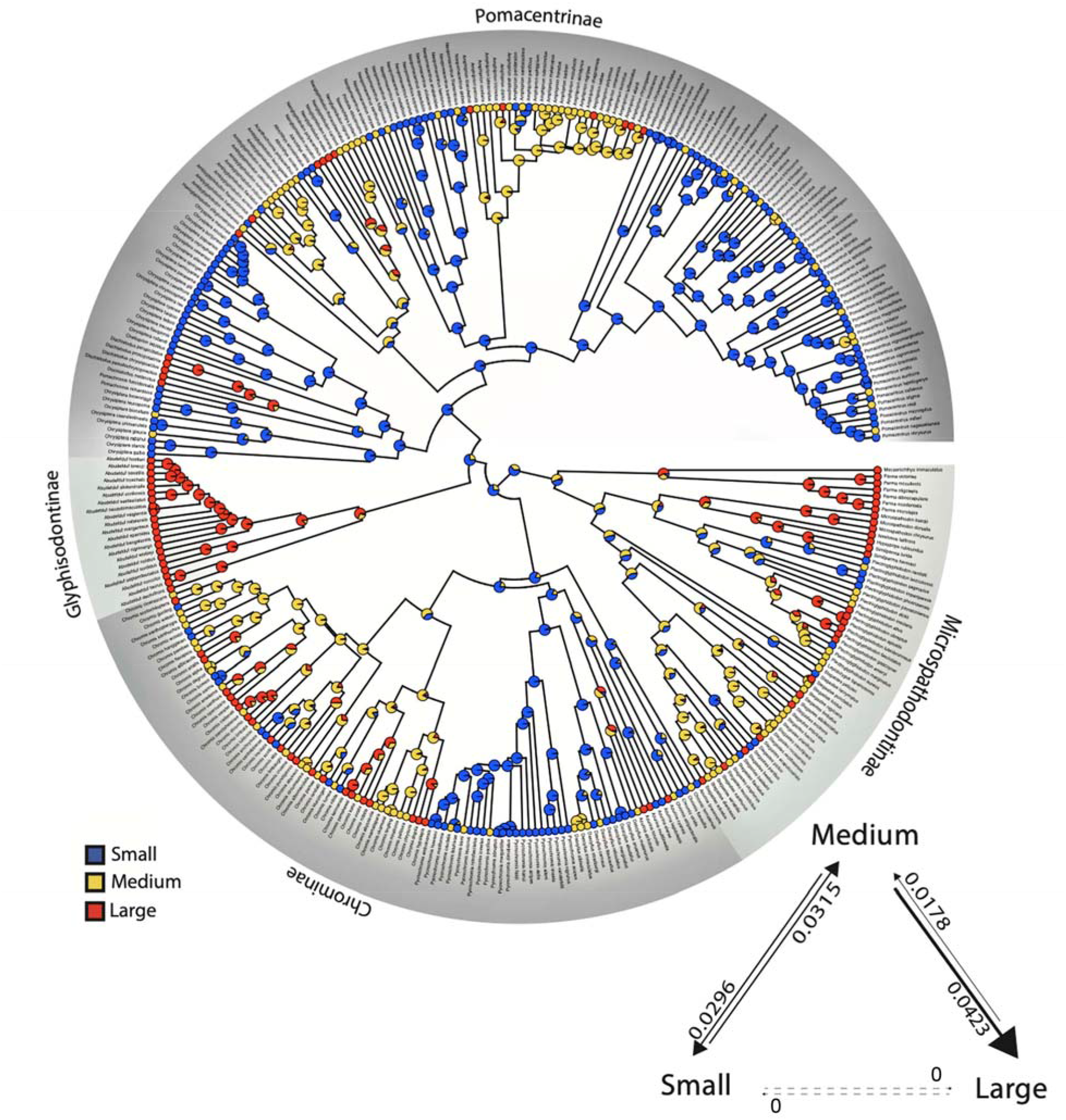
Ancestral state reconstruction of damselfish body size. Maximum body size (total length) gap coded into 3 states with the discretized trait mapped onto the damselfish time-calibrated phylogeny. Transition rates between character states and their rate (if non-zero) between body size states during species diversification throughout the history of the Pomacentridae are shown in lower right. S = small (4.5-10.2 cm TL), M = medium (10.5-14.1 cm TL), L= large (14.4-45 cm TL). Reconstructed state transition counts are S -> M = 46, M -> S = 37, M -> L = 44, L -> M = 15, S -> L = 0, L -> S = 0.

Analysis of the dietary ecotype traits benthic grazer, intermediate omnivore, or pelagic planktivore (Fig. 4) reveals the frequency and asymmetry of transitions between dietary states, with intermediate generalists transitioning to specialist ecotypes with high frequency, particularly among the Pomacentrinae. The Microspathodontinae mostly share a benthic ecotype and there are several other clusters of benthic habits distributed around the tree. Most Chrominae specialize in pelagic feeding (Fig. 4) and this feeding ecotype has evolved repeatedly in damselfish history in large clades (e.g., *Amphiprion* and *Neopomacentrus*) as well as more sparsely in single species, for example in *Pomacentrus* (Fig 4). Ecotype also showed low to moderate levels of phylogenetic signal (λ=0.79, significantly lower than null expectation). We found that transitions between dietary states are prevalent at higher levels in the tree, with complex patterns of evolution and reversal, often involving the intermediate ecotype state among the Glyphisodontinae and Pomacentrinae.

**Figure 4.**
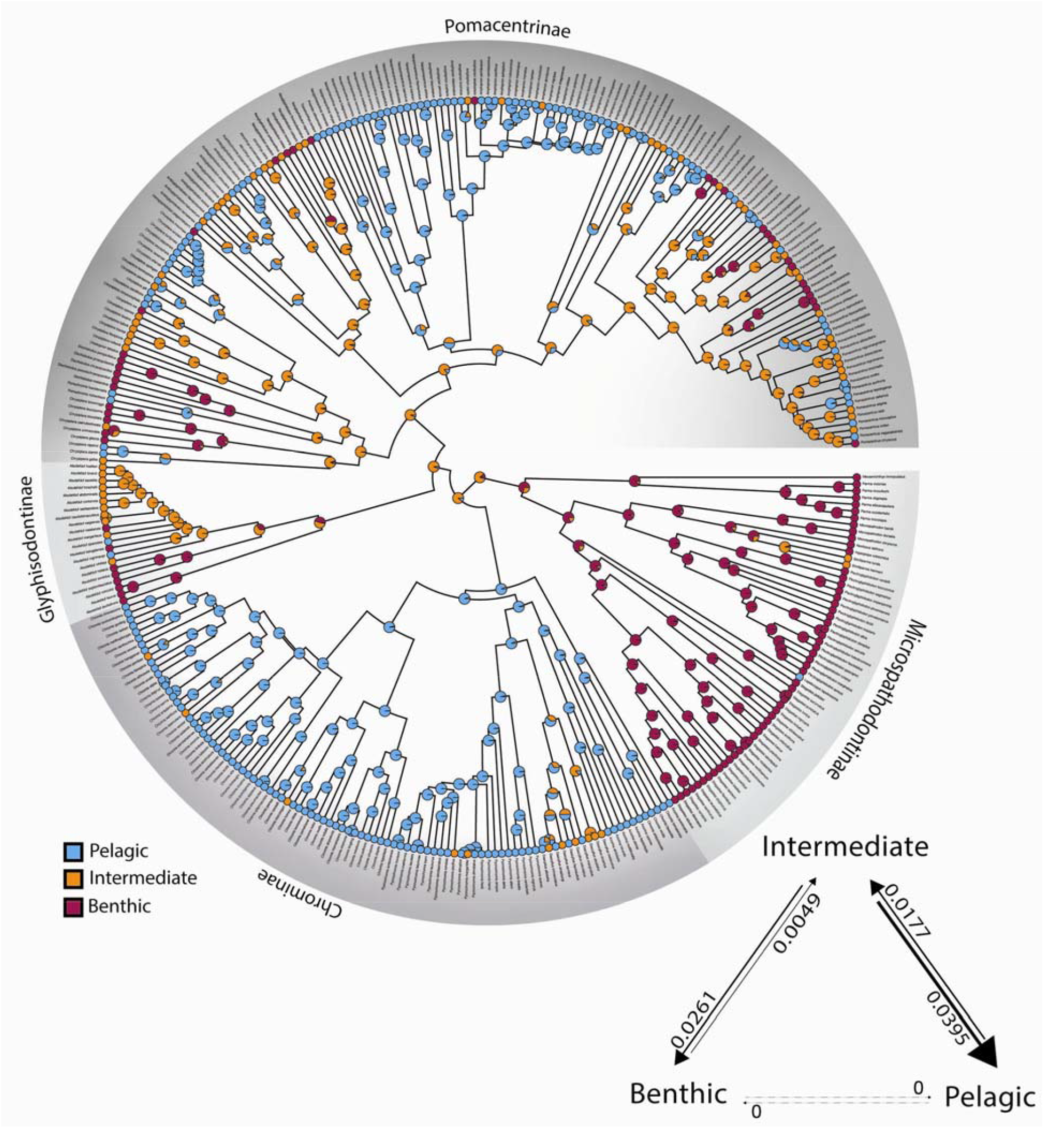
Ancestral state reconstruction of damselfish dietary ecotype. Dietary ecotype traits of benthic, pelagic and intermediate foraging behavior mapped onto the damselfish time-calibrated phylogeny. Transition rates between character states and their rate (if non-zero) between ecotype states during species diversification throughout the history of the Pomacentridae are shown in lower right. Reconstructed state transition counts are: B -> I = 6, I -> B = 25, I -> P = 35, P -> I = 26, B -> P = 0, P -> B = 0.

Algal farming behavior, illustrated in a mirror tree with ecotype (Fig. 5), is restricted to shallow, benthic-feeding species, and is absent from Chrominae and *Abudefduf*. Farming evolved early in damselfish history within the Microspathodontinae as a likely single origin with multiple losses, then again much later, either once or twice in *Chrysiptera*, 3 times in single species of Pomacentrinae 2, and up to 6 times in the genus *Pomacentrus*.

**Figure 5.**
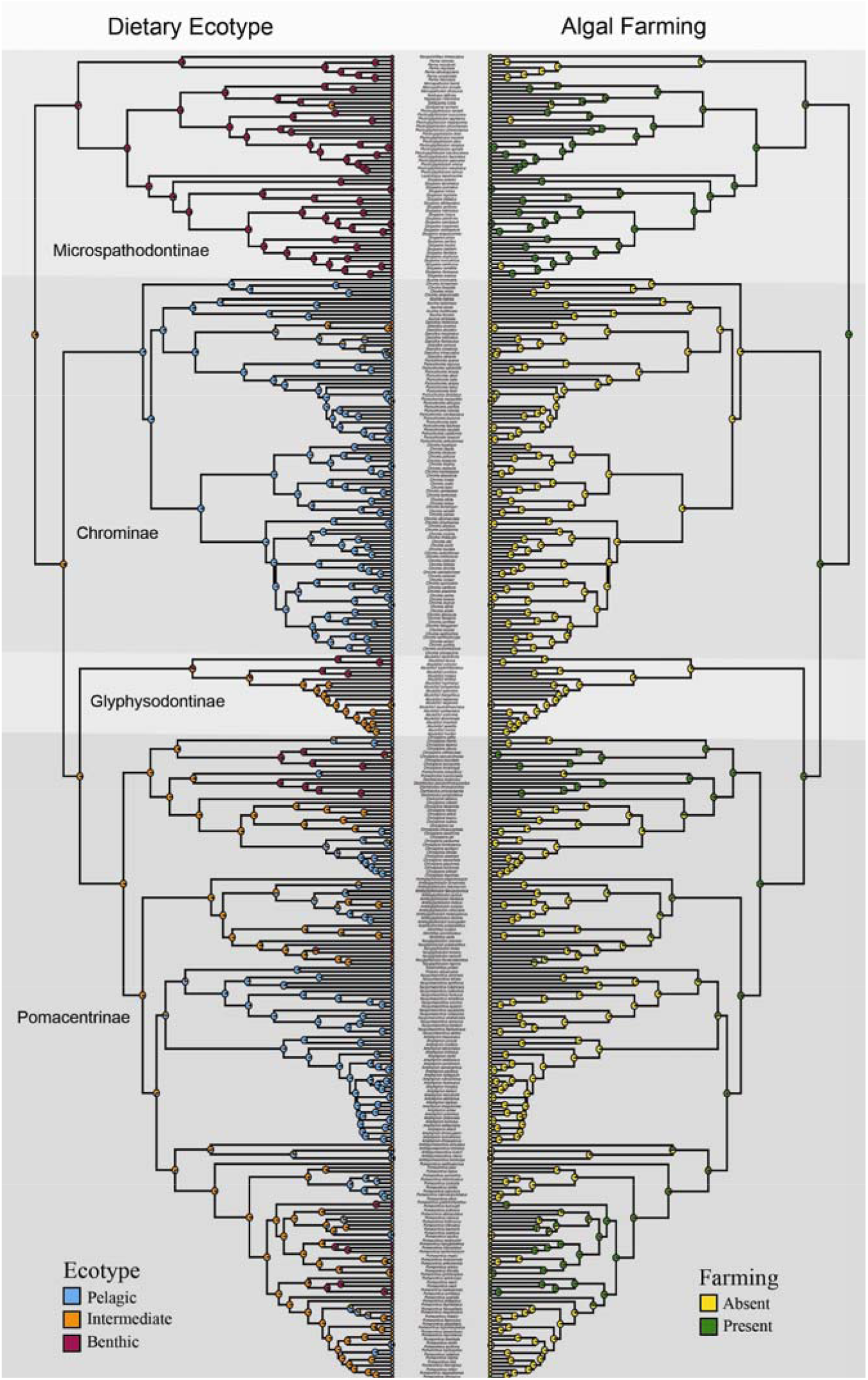
Mirror tree illustration of dietary ecotype and farming behavior. The patterns of dietary ecotype history (left) are illustrated with the presence and absence of the behavioral trait of algal patch “farming” (right) on the damselfish time-calibrated phylogeny.

We tested pairs of traits for character correlations across the entire phylogeny, such as body size (TL) and body depth (BD) associated with ecotype or farming, and none of these tests were significant, suggesting that body size and body depth evolve independently of dietary ecotype and farming behavior. Of particular note is that there is no significant correlation between dietary ecotype and body size. PhylAnova results were: Ecotype vs TL, p = 0.3272; Ecotype vs BD, p = 0.4269; Farm vs TL, p = 0.5062; Farm vs BD, p = 0.9133.

### Diversification patterns of body size and dietary ecotype

The pattern of lineage diversification for the Pomacentridae is that of a highly steady lineage through time plot (Fig. S4), with non-significant difference from a constant rate model when accounting for sampling fraction and without sampling fraction (p=0.73 and p = 0.11, respectively) using the gamma statistic of Pybus and Harvey (71). The results of MuHiSSE models of trait effects on diversification show that both size and ecotype are significantly associated with diversification rate across the phylogeny (Figs. 6, 7). Transitions between states for body size and ecotype are highly asymmetrical (Figs. 3, 4) often being much more frequent in one direction than the other. We ran the transition models under both the assumption that transitions were unconstrained (all pathways equally likely) and that they were constrained to pass through the intermediate state, with similar results. Ecotype transitions are also highly asymmetrical, with the highest estimated transition rate from intermediate-to-pelagic and the lowest was benthic-to-intermediate (Fig. 7).

**Figure 6.**
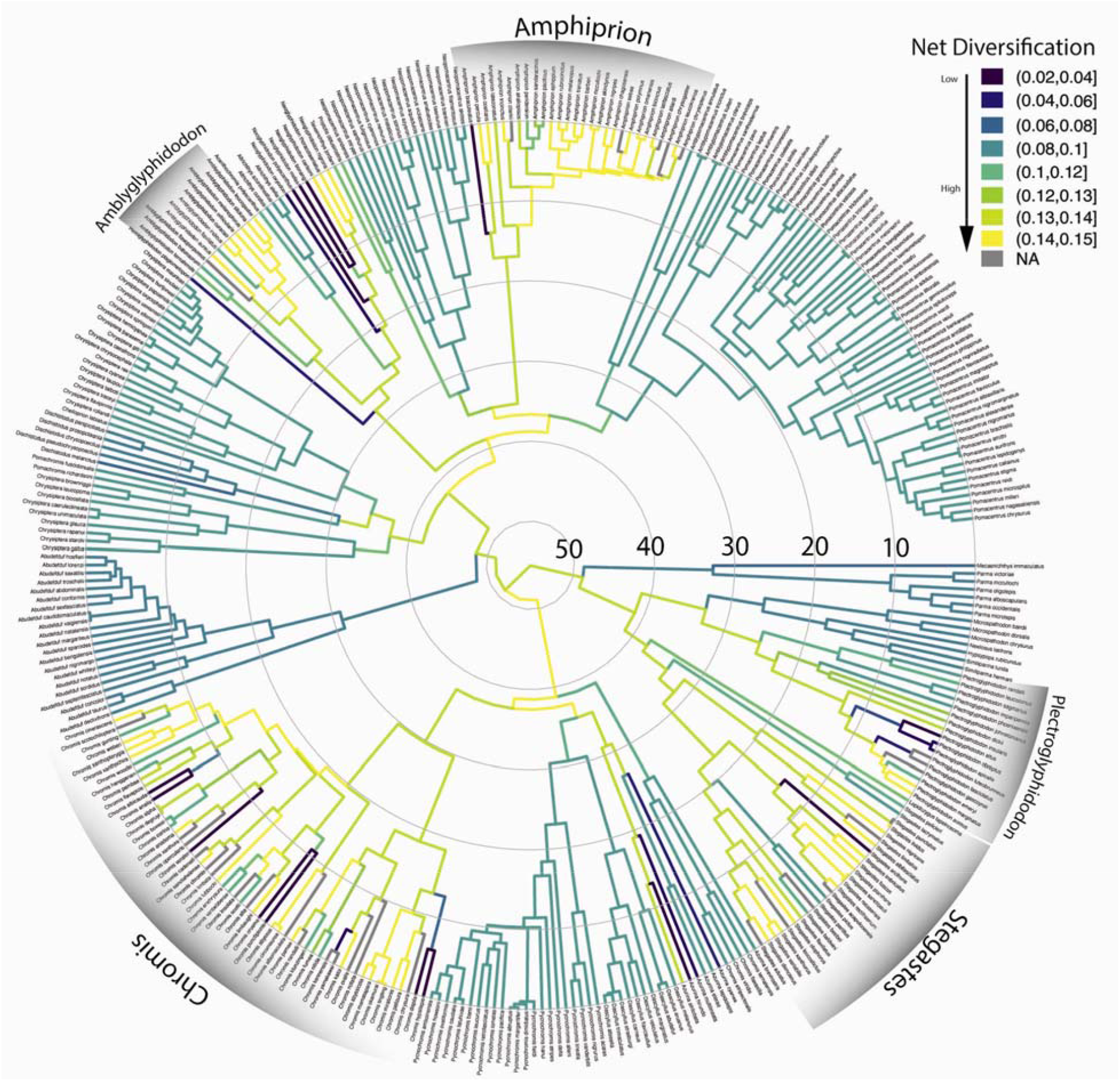
Body size diversification rates on the time-calibrated phylogeny of the Pomacentridae. Body size shows multiple areas of elevated diversification in lighter colors throughout the tree *(Amblyglyphidodon, Amphiprion, Chromis, Plectroglyphidodon and Stegastes)*.

**Figure 7.**
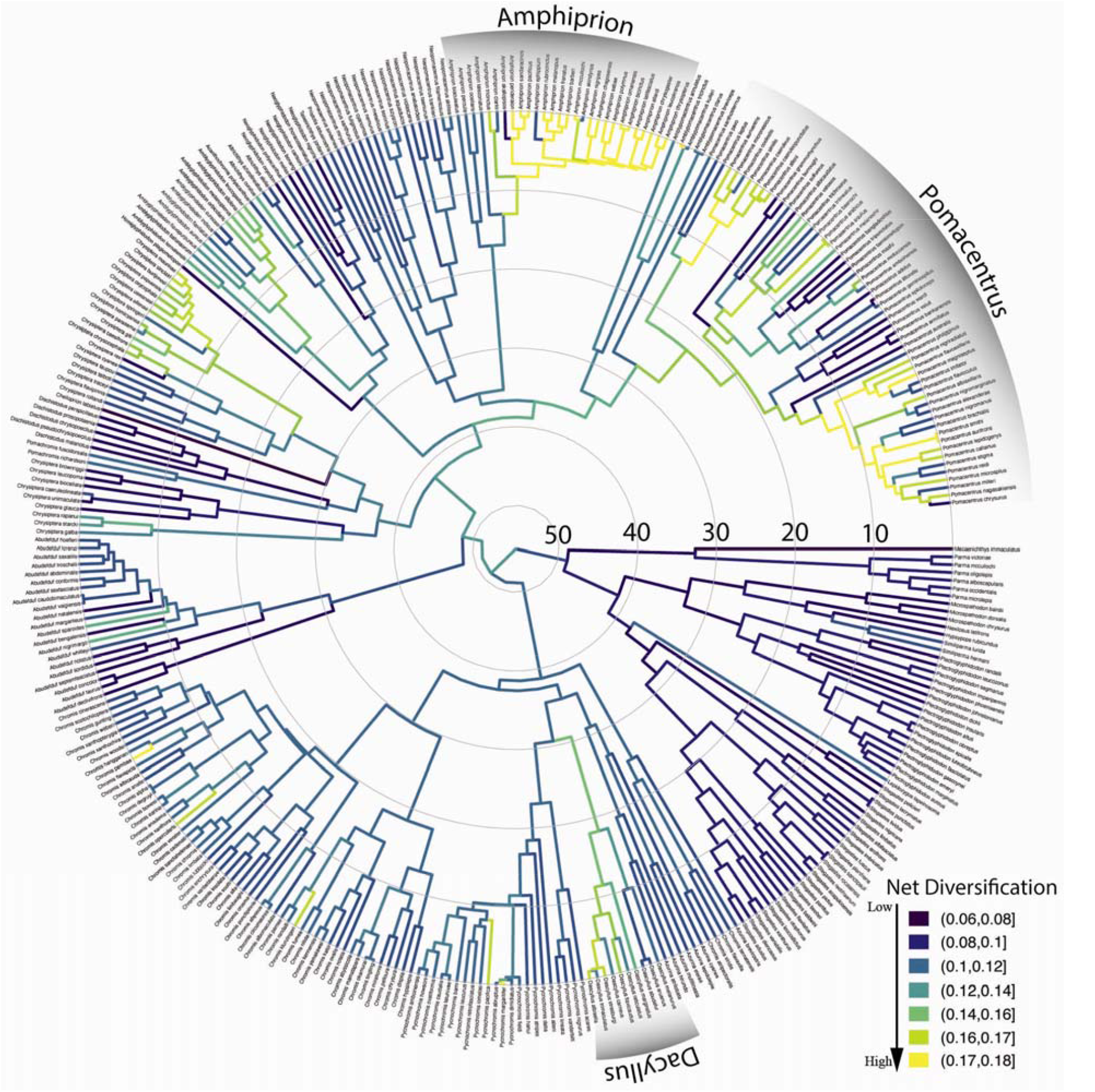
Ecotype diversification rates on the time-calibrated phylogeny of the Pomacentridae. The relatively low, steady pattern of diversification in the dietary ecotype trait is punctuated with several independent origins of elevated ecotypic diversification rate (*Amphiprion* is the most prominent) in lighter colors throughout the tree.

Size-dependent models that include hidden states were strongly favored, with the best supported model being MuHiSSE relaxed (ωAIC = 0.57), followed by Large Absorbing (ωAIC = 0.42), with other models having much lower AIC scores (Table 1). Diversification rates were highest among medium sized damselfishes, followed by small and then large categories (Fig. 6, Fig. 8A). Ecotype-dependent models that include hidden states were also strongly favored, with the best supported model being MuHiSSE (ωAIC = 0.46), followed by MuHiSSE relaxed (ωAIC = 0.32) and Benthic Absorbing (ωAIC = 0.18) (Table1). These models differ in the parameterization of transitions among hidden states and transitions out of the benthic state. All other models had ωAIC values < 1%. When averaged across all models and over hidden state reconstructions, mean net turnover and net diversification were estimated to be highest in the pelagic state, followed by intermediate and then the benthic state (Fig. 8B). These higher rates of diversification are especially prevalent in the Amphiprionini, several parts of the *Pomacentrus* tree, and in *Dascyllus* (Fig 6). The particularly elevated diversification distribution among pelagic damselfishes seen in Fig. 7 represents the anemonefishes.

**Figure 8.**
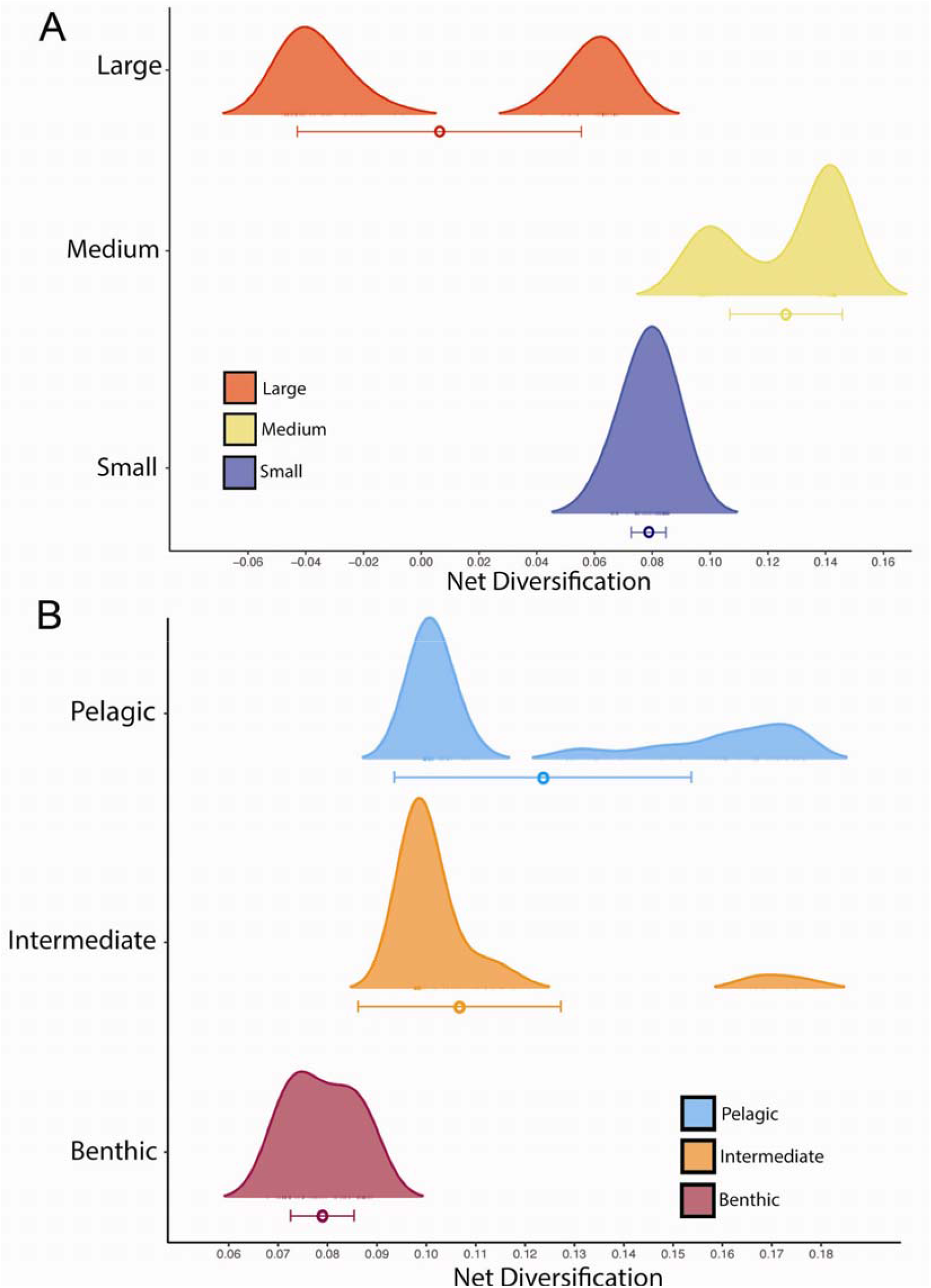
The distribution of net diversification across (A) damselfish size classes, and (B) damselfish ecotypes. Model averaged mean and standard deviation of diversification for each size class and ecotype is indicated by a colored circle and bar, respectively. The height of the peak indicates the number of tips for each ecotype with that particular estimated net diversification rate. Mean estimated net diversification was highest in the medium size class and the pelagic ecotype state.

## Discussion

The synthesis of species-rich phylogenies and diverse morphological, functional, and ecological trait data yields new ways of visualizing evolutionary patterns, promotes the development of novel hypotheses, and empowers their testing using comparative methods. Marine fishes offer particularly revealing opportunities for these explorations due to the recent development of corresponding phylogenies and datasets (organismal and/or ecological characters) for species-rich lineages (19,21). Integrative phylogenetic approaches to reef fishes allow us to address important questions associated with local and global biogeographic patterns, the history of structural and functional evolution, and the diversification of important traits that have driven species richness for high diversity reef fish families. The central conclusions of this work are that (1) the core structure of the damselfish phylogeny is resolved with high support, with intriguing questions remaining to be answered using larger datasets, (2) multistate hidden-state speciation and extinction models show that transitions between body sizes and between dietary ecotype states are a significant influence on damselfish diversification, (3) there is pronounced asymmetry in transition directionality between character traits in body size and ecotype that is a significant driver of diversification patterns in damselfishes, and (4) the convergent pattern of ecological diversification that has been described as a characteristic of the Pomacentridae as a whole is limited mostly to the Indo-West Pacific subfamily Pomacentrinae.

### Damselfish phylogeny: robust clade support and time-tree resolution

Our understanding of phylogenetic relationships among the Pomacentridae has improved tremendously in completeness, resolution and support as species sampling and character matrix density has increased. From early work by Tang (34), Jang-Liaw (31) and Quenouille et al. (33) to Cooper et al. (1), Frédérich et al. (15), Tang et al. (37) and the present study, the molecular phylogeny of damselfishes has grown from 23 species to now include 345 of the 422 known species. All regions of the tree are densely sampled, and although many species are represented by just one or a few gene sequences, the overall topology is well-resolved and generally well-supported under both likelihood and Bayesian approaches.

The recent phylogenetic analysis and much-needed taxonomic treatment of the Pomacentridae using 8 sequence loci and 322 damselfishes by Tang et al. (37) significantly revised pomacentrid taxonomic names and enhanced our understanding of damselfish relationships. Their study, released after the preprint and during review of the present study (52), allowed us to adopt the new taxonomic framework and compare our analysis of 12 loci for 345 damselfishes to this nearly simultaneous work. The central conclusion from this comparison is that there is extensive agreement between the two studies, with some notable exceptions, detailed herein.

The phylogeny of the subfamily Microspathodontinae (Fig. 1) has been relatively stable over the past decade. Taxonomic issues have largely been resolved (37,70), but other puzzles remain. *Parma* species are resolved as the monophyletic sister group to *Mecaenichthys*, as we have eliminated the older sequence data for *Parma oligolepis* that appears to be a misidentification or contamination linking it to *Stegastes. Nexilosus* is now resolved as sister to *Hypsypops*, using recent CO1 data only, and eliminating the older sequences for this species that were generated using ancient DNA approaches on a preserved specimen (1). However, the resolution of *Nexilosus* in this position remains uncertain and its relationship to *Similiparma* merits further investigation. *Lepidozygus* is sister to *Stegastes*, with that clade sister to a monophyletic *Plectroglyphidodon* Although the Bayesian posterior support for *Lepidozygus* in this position is high, we note that the long branch for this species has made it hop around in recent analyses (1,15,33) so view its position here with caution. These relationships are in agreement at both deep and finer levels with the Tang et al. (37) topology.

Our analysis resolved the next major group up the damselfish tree to be the subfamily Chrominae, with high support in all analyses. This is strikingly different than the recent Tang et al. phylogeny, which resolved *Abudefduf* in this position. The overall topology of the pomacentrid subfamily Chrominae (Fig. 1) has been revised considerably across recent phylogenetic hypotheses, with several solidly monophyletic groups of taxa that typically render *Chromis* itself polyphyletic. *Chromis* is rendered polyphyletic in our analysis (in contrast to Tang et al. (37)) by a group of 4 species (*C. atripectoralis, C. favaxilla, C. ternatensis*, and *C. viridis*) forming a strongly supported sister group to the rest of the subfamily (Fig. 1). The genus *Azurina* (now including several species previously in *Chromis*) is resolved as monophyletic, though with weak support, forming the sister to *Dascyllus* plus *Pycnochromis*. However, we also show that the finer, detailed species relationships within the Chrominae are uncertain, with many clades of *Chromis* and *Pycnochromis* poorly supported (Fig. 1), so we suggest that detailed evolutionary studies focused on within-chromine patterns require additional research.

The phylogenetic tree resolved here places the Glyphysodontinae (genus *Abudefduf*) as the sister to the crown Pomacentrinae, with high support. This is a major difference between our topology and that of Tang et al. 2021, which may due to our increased species sampling, the greater number of genetic loci sampled, or both. We anticipate that still greater sampling using phylogenomic approaches may resolve this discrepancy. Recent phylogenetic work (1,15) and analyses focused on the genus *Abudefduf* (72,73) have provided both Sanger and UCE sequence data for all 21 known species, rendering *Abudefduf* as one of the most clearly resolved, monophyletic pomacentrid groups (Glyphysodontinae; Fig. 2). The time calibrated reconstruction here of about 30 mya for the genus is slightly older than the 25 mya recovered by Campbell et al. (73), due to new time-calibrated outgroup nodes in the present tree (60). However, the 25 mya age remains within the 95% highest posterior density error bars of our estimate. The species-level topology of *Abudefduf* shown here is also similar but not identical to that resolved under ML for ultra-conserved element data (73), with the positions of *A. concolor* and *A. lorenzi* being the primary differences, and we suggest that the topology based on the larger UCE dataset may be preferred.

The subfamily Pomacentrinae is full of phylogenetic intrigue, taxonomic issues, and evolutionary fascination (Fig. 2). Pomacentrinae 1 contains the two main clades of *Chrysiptera* with a clade of *Pomachromis* + *Dischistodus* (sister genera, each resolved as strongly monophyletic) nested between them with high Bayesian posterior probability, and support in other analyses as well (Fig. S1, S2). The relationships among *Chrysiptera* species resolved here largely agree with recent analyses of the genus (37,74,75), although the taxonomy of *Chrysiptera* within Pomacentrinae 1 is the last large damselfish group likely to require additional significant taxonomic revision. Pomacentrinae 2 (the Hemiglyphidodontini) is a diverse group of damselfishes whose relationships are congruent with current taxonomy, with *Hemiglyphidodon, Amblyglyphidodon, Acanthochromis, Altrichthys*, and *Neoglyphidodon* all resolved as monophyletic. Pomacentrinae 3 (Fig. 2) is resolved as an unnamed group with high support (perhaps requiring tribe status), containing *Pristotis, Teixeirichthys*, and *Neopomacentrus* as the sister group to the clownfishes. This topology is recovered with high support in our analyses, but is significantly different from the Tang et al. topology, in which this clade was recovered as sister to *Pomacentrus*.

The remarkable clade of anemonefishes, or clownfishes (Tribe Amphiprionini; Pomacentrinae 4; Fig. 2) is resolved with high support as a monophyletic clade that diversified almost 15 million years ago and underwent a burst of diversification during the past 5 million years. These findings are consistent with other recent estimates of 5-13 mya for the origin and radiation of the Amphiprionini (15,76,77). Litsios et al. (76) supported the long-held hypothesis that mutualism with sea anemones promoted the adaptive diversification of the anemonefishes. Marcionetti et al. (78) identified 17 genes that underwent positive selection in the early stages of the anemonefish radiation, two of which are potentially associated with chemical components of the toxin discharge mechanisms of sea anemones. The anemonefishes are mostly of medium size (Fig 3), with the peculiar trait of being closely tied to their benthic anemones but mostly feeding above it, in the water column, on zooplankton (Fig. 4). There are a number of transitions among ecotypes and among size classes in the clade, which combined with their rapid diversification make them an important part of the trait diversification history of the family discussed below.

Our results show strong support for the monophyly of *Pomacentrus* and the sister-group relationship between *Amblypomacentrus* and *Pomacentrus* (Pomacentrine 5; Fig. 2), which is the second largest genus of damselfishes, and good support for most of the relationships within this clade (Fig. 2). However, it should be noted that most of the resolution within the *Pomacentrus* tree is supported by barcode or other small loci such as d-loop, provided by recent new species descriptions (79,80). This phylogeny for *Pomacentrus*, which contains 58 of the 80 recognized species, is another step forward in the species tree of this genus, as prior phylogenetic work has included up to 34 *Pomacentrus* species (15) or more recently 52 species (37).

In contrast to the other major clades, the Pomacentrinae are restricted to the Indo-West Pacific and the majority of them inhabit coral reefs (1,70). Although no extant pomacentrine damselfishes are native to either the Atlantic or East Pacific, their late Eocene divergence date (42 mya) considerably precedes the closing of the circumtropical Tethys seaway between 12 and 18 mya (Early Miocene; Fig. 2) when warm-water connections between the Atlantic and Indo-West Pacific were severed (81). Their diversification across the tropical Indo-West Pacific and their absence on coral reefs of the Atlantic/East Pacific is therefore somewhat of a paradox.

Most major branching events in pomacentrine evolution roughly coincide with the Eocene-Oligocene Transition about 33.5 mya (Fig 2), a significant global biotic reorganization with major changes to ocean ecosystems (82), suggesting that these fishes may have benefitted from this period of ecological upset. Yet the steady radiation of pomacentrines throughout the tropical Western Pacific is in contrast to their absence from the Eastern Pacific, which may be the result of a coral extinction event in the late Oligocene and Early Miocene. Between 24 and 16 mya, a large portion of the reef-building coral genera that inhabited what is now the Caribbean went locally extinct, but persisted in the Indo-West Pacific (83,84). This loss of Caribbean coral reef habitat extended into the period during which the Tethys Seaway closed 12-18 mya (85). Coral reef habitat loss in the Caribbean may have caused pomacentrine extinctions, and the Tethyan closure, the East Pacific Barrier, and colder ocean waters around Southern Africa prevented their recolonization of the Atlantic and Eastern Pacific. The damselfishes are a rich system for more detailed exploration of biogeographic patterns and reef fish community assembly across the marine realm.

### Trait transitions and asymmetric damselfish diversification patterns

The damselfishes are a curious example of a lineage that has undergone a steady global diversification, ultimately resulting in a somewhat constrained adaptive radiation (14,15). Damselfish diversification to become one of the largest and most important clades on coral reefs is characterized, and perhaps driven by, complex patterns of divergence and convergence in size, shape and feeding ecology, within a curiously bounded set of ecomorphological end points. The regular, clock-like nature of damselfish speciation through time (Fig. S4) rejects early-burst or late-burst evolutionary hypotheses (86) for the generation of high species richness. The metronomic pace of damselfish radiation (excepting the recent speciation of *Amphiprion*) is in contrast to patterns seen in some fish groups such as parrotfishes (13,16), cichlids (87) and goatfishes (88). These steady diversification rates are significantly associated with both body size and ecotype transitions in these fishes, in combination with one or more hidden (non-focal) states or traits according to our MuHiSSE tests of trait effects on diversification across the phylogeny (Figs. 6, 7).

The evolution of body size in damselfishes is strongly associated with their diversification, via two quite different phylogenetic patterns: frequent change back and forth among states in some clades or getting largely stuck on a single state in other clades. The best-fit model for the influence of size on damselfish diversification was the MuHiSSE relaxed state-dependent model with a hidden state, although a close second in model fit was the large-absorbing model (Table 1). Elevated levels of size diversification occur primarily in clades with a mosaic pattern of all three size states (Fig. 3) and a high prevalence of intermediate size transitions to either small or large species, such as in *Chromis, Stegastes*, and *Amphiprion* (Fig. 6). Supporting the large-absorbing model, there are several clades with large body size that evolved the trait at or near their origin and maintained it during moderate levels of diversification, including the microspathodontine giant damselfishes, *Abudefduf* and *Dischistodus* (Figs. 3, 6). Frequent two-way transitions between small and medium sizes, accompanied by largely one-way medium to large transitions with subsequent speciation in some clades have shaped the history of the pomacentrids and are clearly detected by hidden state modeling.

Episodes of evolutionary diversification in size are common at multiple levels in fishes, often alternating with patterns showing stability in size across clades or across time intervals. This has been illustrated by broad-scale analysis of the size of fossil fishes across 500 million years (89), exploration of the association of body size with habitat or trophic position at multiple scales across living fishes (90–93), and examining the evolution of size among particular clades of marine and freshwater fishes (19,94,95). The phenomenon of directional evolution toward large size has been particularly well studied in Clupeiformes (95) and in migratory fishes more broadly (94) where large body size evolves among migratory clades, a system in which larger size confers clear advantages for swimming endurance. Among Clupeiformes (95) and at broader phylogenetic levels (94), migratory clades often appear to get relatively stuck on the large size trait, with few reversions, a pattern similar to that found here for damselfishes. Larger body sizes are also associated with increased gonad size and reproductive output (96,97), which is a possible driver of the strong large-absorbing model performance in the Pomacentridae, as migration is not thought to occur to any meaningful degree in damselfishes. Large body size in damselfish groups such as *Dischistodus* have been shown to be associated with particular skull morphology associated with feeding (98), and the association between body size and feeding has often been cited as being significant across the family (19). However there is no significant relationship between dietary ecotype and body size in damselfishes, with a highly non-significant PhylANOVA p-value of 0.32.

The history of multistate transitions among dietary ecotypes was also significantly associated with pomacentrid diversification, with the best-fit model the MuHiSSE state-dependent model with a hidden state, MuHiSSE relaxed a close second, and benthic-absorbing third (Table 1). These results show that both body size and ecotype are strongly associated with pomacentrid diversification throughout their history, although there are certainly other traits, perhaps the detected hidden trait, likely to be important influences on diversification, from anatomy, reproductive biology and color pattern to behaviors such as territoriality and parental care. Similar to the evolutionary pattern for body size, transitions among dietary ecotypes in the damselfishes are strongly asymmetrical (Figs. 4, 7), with transitions through intermediate states the rule, frequent back-and-forth transitions in some groups, but with some clades often getting somewhat evolutionarily “stuck” at the extremes. This dual pattern of ecotype evolution is clear (Fig. 4) with the microspathodontines largely stuck on benthic (purple), the chromines almost uniformly pelagic aqua, and the rest of the family presenting a more diverse and nuanced ecotypic evolutionary pattern. The relatively high score of the Benthic Absorbing model in the MuHiSSE analysis (Table 1: 3^rd^ place) reflects this tendency of some damselfish groups to go benthic and diversify in that state, apparently without much reversal. The original Nakov et al. (50) development of MuHiSSE approaches tested Absorbing models of diversification, for freshwater and marine absorbing hypotheses for diatom diversification, but found little support for them. To our knowledge, others have not tested this idea that a trait in stasis (the absorbing model) may be a significant correlate of diversification, and here for damselfishes we find that the absorbing models are strongly supported for large body size and moderately supported for benthic dietary ecotype.

It is intriguing that the Pomacentridae exhibit just three major feeding niches: planktivory, omnivory and herbivory. Diversification of the damselfishes has not involved expansion into an increasing number of ecological niches, but rather repeated convergence on a limited set of similar feeding morphologies and ecologies. This pattern has been described as driving damselfish diversification due to the repeated pattern of convergence on these three primary ecotypes (14), and has been considered to be characteristic of the damselfishes as a whole.

However, the present analysis of dietary ecotype using a more comprehensive phylogeny shows that it is primarily the Pomacentrinae that have undergone this ecotype diversification (Figs. 4, 7). Although all three ecotypes are distributed across the phylogeny, most major clades specialize on a single ecotype, or exhibit a small number of ecotype transitions (Fig. 4, 5). Examples of this include the typically large body size and herbivorous ecotype of the microspathodontine clade. At the other extreme, almost all Chrominae are relatively small and planktivorous, with the *Dascyllus* lineage departing from this pattern as a rare reversion to the intermediate state within the clade (Fig. 4, 5). Although most *Dascyllus* species feed on plankton, many also consume considerable amounts of algae (99–101). Their trophic morphology is characteristic of omnivorous damselfishes and the feeding kinematics of *Dascyllus aruanus* are similar to those of other pomacentrid omnivores (39). The Glyphisodontinae (*Abudefduf*) are a relatively species-poor lineage but they have evolved all three ecotypes, with only a small amount of the back-and-forth pattern in dietary and body size transitions seen among the Pomacentrinae (Fig. 4).

The Pomacentrinae (and in particular *Pomacentrus*) show a much stronger pattern of ecotypic convergent evolution, with repeated transitions between all three ecotypes (Figs. 4, 7). Although the Pomacentrinae exhibit the highest degree of geographic and habitat limitations of any of the subfamilies (only found in the Indo-West Pacific; almost entirely restricted to coral reefs) and are one of the two most recently evolved major clades, they have produced roughly twice as many species as the Chrominae, almost three times as many species as the Microspathodontinae, and are approximately ten times more species-rich than their sister lineage Glyphisodontinae. Members of all five damselfish subfamilies can be found on Indo-West Pacific coral reefs, but only the Pomacentrinae have demonstrated a pronounced ability to frequently transition between three ecotypes that lie along a bentho-pelagic axis within these ecosystems. The enhanced evolvability of the Pomacentrinae relative to other damselfishes may have contributed to the tremendous success of this lineage.

Algal farming or gardening behavior is a complex behavioral trait that has evolved multiple times in four main damselfish lineages, occurring in fishes of diverse body sizes but restricted to the shallow benthic habitats where desirable algae grow (Fig. 5). Farming in the Micropathodontini probably evolved once near the base of the clade, with 9 subsequent losses, although alternative character reconstructions involve up to 9 independent origins in this group (Fig. 5). The Cheiloprionini likely evolved farming once with 4 losses, the Pomacentrinae 3 group twice in *Hemiglyphidodon* and *Neoglyphidodon*, and up to 6 origins in the Pomacentrinae (or two origins with multiple losses). Cultivating and defending an algal patch has long been recognized as an unusual level of behavioral complexity in fish feeding behavior (42,102,103), the intricacy of which recently increased as it has been shown to include the domestication of mysid shrimps by damselfishes to help fertilize the garden (104). All damselfish farmers are benthic, yet most benthic feeders are not farmers, and there is no significant character association between farming presence/absence and dietary ecotype across the phylogeny. The independent origin of this complex behavior up to a dozen times across the tree, associated with intense aggressive territorial defense behavior, and associated each time with the benthic ecotype that shows the lowest diversification rate across the tree, makes this a promising system for study of the factors that accelerate or slow diversification in damselfishes.

### Time tree structure and species sampling are critical in comparative analyses

A central conclusion of this study is that developing and using a nearly complete tree topology for the damselfishes, with extensive data matrix curation, high species richness across clades and updated time calibrations, can substantially change our interpretations of evolutionary patterns, trait associations, and the factors driving diversification. In addition to changes in our perspective on the ages of major groups and our understanding of the influences of body size and ecotype on diversification, the present work challenges previous conclusions about the importance of herbivory in reef fish diversification and the association of body size with diet.

The time-calibrated phylogeny resolved here sets the age of the damselfish split from other related families at about 80 million years ago, and the root age of the first split within damselfishes to 56 mya (Fig.1, Fig. 2, Fig. S3). This basal node age of the pomacentrids at 56 mya agrees with two previous estimates of the time tree for damselfishes alone (105) and within the broader fish phylogeny (93), and is close to the Frédérich et al. (15) time-tree result of 52 mya. However, most other studies placing pomacentrids within larger phylogenetic contexts among have derived much younger ages for the family including 25 mya (106), 36 my (60,107), and 48.7 mya (108). The comprehensive online time-tree for fishes in this latter study (108,109) including just 223 damselfish species and resolving a 12% younger age, is often harvested for comparative analyses involving damselfishes. While acknowledging the value of open access sources for phylogenetic trees, comparative analyses using the pomacentrid tree therein may be misleading. Not that the present tree is necessarily “correct”; it is likely that the root age of damselfishes will be pushed back yet deeper in time, as most large phylogenomic datasets for fishes are trending toward older family ages. However, the revised time-tree and increased topological richness of the current phylogeny for the family may benefit online tree-of-life resources and future comparative work requiring a species-rich phylogeny.

Trait analysis using the current tree for damselfishes leads to conclusions about the history of diversification, and its association with key reef fish traits, that differ from prior work. Prior trait analyses using phylogenetic trees of approximately 200 species (19) concluded that body size and dietary ecotype are correlated throughout damselfish history, whereas our analysis of these traits for 345 species lead us to conclude that this is not the case-neither phylogenetic nor raw-data correlations between size and ecotype are significant. Prior work that was the first to apply SSE modeling to the damselfishes (38) showed high rates of transitions out of the intermediate dietary ecotype into the pelagic and benthic ecotypes, similar to the present analysis using multi-state rather than binary models. However, in contrast to this prior work (38), which concluded that the intermediate ecotype has the highest diversification rate, we found higher diversification rates associated with the pelagic ecotype, followed by the intermediate and benthic ecotypes. This is likely due to the present study having a larger, more complete sample of the tree and the traits, and we anticipate that some of the present findings may be similarly revised once the entire family is resolved. Finally, a recent multi-family reef fish analysis (21) drew the general conclusion that clades with larger body sizes and herbivorous clades have the highest diversification rates, and yet the damselfishes, one of the largest reef fish families, show exactly the opposite results. Damselfishes may simply be different, perhaps due to their tendency toward trophic territoriality and/or their unusual (for reef fishes) nesting and egg brooding behavior. We agree with the recent suggestion by Miller et al. (110) that large, multi-clade analyses benefit from sub-clade explorations, as a combination of family-specific analysis with larger multi-clade patterns will be necessary to understand the diversification of reef fishes.

In summary, an expanded, time-calibrated damselfish phylogeny allows us to make important additions to our understanding of damselfish ecological radiation. Body size and dietary ecotype are both associated with damselfish diversification, with strong asymmetry in the direction of body size evolution and ecotype transition, with shifts away from the intermediate omnivorous ecotype to either the pelagic or benthic ecotype. It is primarily the crown group in the family, the Pomacentrinae, showing the strongest pattern of convergence in ecotype. All three ecotypes have evolved repeatedly across the damselfish phylogeny, but before the appearance of the Pomacentrinae, transitions in ecomorphology were more likely to be associated with major branching events rather than at finer levels within clades. Increased phylogenetic resolution, feeding mechanics studies, dietary ecology and developmental biology of the damselfishes will continue to reveal surprising evolutionary trends in this spectacular group of fishes.

## Acknowledgments

Thanks to Lydia Smith and Jillian Henss for assistance with DNA sequencing, and to the Division of Fishes and the Pritzker Molecular Laboratory at the Field Museum of Natural History for collections and data acquisition support. We also thank the fish collections of the Australian Museum, Scripps Institution of Oceanography, National Museum of Natural History in the Smithsonian Institution, Institute of Zoology in the Academia Sinica, and the Gil Rosenthal lab of Texas A & M University for supplying pomacentrid tissues. Supported by NSF grant DEB 1541547.

## Supplemental Data

https://figshare.com/articles/dataset/Supplemental_Data_Revised_Phylogeny_of_the_Damselfishes_Pomacentridae_and_Patterns_of_Asymmetrical_Diversification_in_Body_Size_and_Feeding_Ecology/16663633

## Supporting information

Text S1. Supplemental Methods and Results.

FigS1. Best ML tree. Maximum likelihood topology for the damselfishes.

FigS2. Time tree posterior support. Time-calibrated tree for the damselfishes showing posterior probability support for nodes of the tree.

FigS3. Time tree node ages. Time-calibrated tree for the damselfishes showing reconstructed ages of nodes of the tree.

FigS4. Lineage through time plot for the damselfishes.

TableS1. Genbank accession numbers. Genbank accession numbers for the 12 genes and 350 taxa used in the damselfish phylogenetic analysis, with color coding by major data contributor.

TableS2. Primers. A list of primers used in gene sequencing.

TableS3. Trait data for damselfishes. Trait data on ecotype, farming behavior, body size, and body depth for the Pomacentridae.

TableS4. Ecotype and diet for damselfishes with reference notess.

TreeFile1 DamselBestML.tre. Treefile for the maximum likelihood topology of the damselfishes.

TreeFile2 DamselTimeTree.tre. Treefile for the time-calibrated topology of the damselfishes from BEAST analysis, with 5 outgroups.

TreeFile3 DamselsOnlyTimeTree.phy. Treefile for the time-calibrated topology of the damselfishes from BEAST analysis, damselfish only, in .phy format for upload in R.

